# HPV and p53 status as precision determinants of head and neck cancer response to DNA-PKcs inhibition in combination with irradiation

**DOI:** 10.1101/2023.11.02.565300

**Authors:** Liana Hayrapetyan, Selina M. Roth, Lusine Hovhannisyan, Matúš Medo, Aurélie Quintin, Julien Ott, Joachim Albers, Daniel M. Aebersold, Yitzhak Zimmer, Michaela Medová

**Affiliations:** Department of Radiation Oncology, Inselspital, Bern University Hospital; Department for BioMedical Research, Radiation Oncology, University of Bern; Graduate School for Cellular and Biomedical Sciences, University of Bern; Research Unit Oncology, the healthcare business of Merck KGaA, Darmstadt, Germany

## Abstract

Major risk factors of head and neck squamous cell carcinoma (HNSCC) are tobacco use and human papillomavirus (HPV). HPV E6 oncoprotein leads to the degradation of the p53 protein, whereas HPV-negative cancers are frequently associated with TP53 mutations. Peposertib is a potent and selective, orally administered small-molecule inhibitor of the catalytic subunit of the DNA-dependent kinase (DNA-PKcs), a key regulator of non-homologous end joining (NHEJ). NHEJ inhibition along with irradiation (IR)-induced DNA double-strand breaks has the potential to increase antitumor treatment efficacy. Here, we investigated the responses of HNSCC models with distinct HPV and p53 status to treatments with IR, DNA-PKcs inhibition, and their combination.

We observed that IR-induced DNA damage combined with peposertib administration shortly before IR results in decreased cell viability and proliferation and causes DNA repair delay in all the studied HNSCC cell lines. However, our data confirm that the actual cell fate upon this treatment is strongly dependent on cellular p53/HPV status. Cells lacking functional p53 due to its degradation by HPV or due to the presence of a loss-of-function mutation are arrested in the G2 phase of the cell cycle and eliminated by apoptosis whereas p53-proficient HNSCC cell lines undergo senescence. Consequently, HPV+ cancer cell lines and xenografts display stronger and more durable responses and seem to benefit from the combined treatment more than p53-proficient HNSCCs. In conclusion, DNA-PKcs inhibitor peposertib should be further studied as a potential radiosensitizer for HNSCCs, taking into consideration the genetic background and the HPV status of a particular tumor.

## Introduction

Head and neck squamous cell carcinomas (HNSCCs) are among the most frequently diagnosed tumors worldwide, with mortality exceeding 500,000 deaths yearly [Sung et al, 2021]. Major risk factors leading to the development of this malignancy include various environmental substances and pollutants (e.g., tobacco use, excessive alcohol consumption) as well as chronic infection with cancerogenic strains of human papillomavirus (HPV) and Epstein-Barr virus [Johnson et al, 2020]. While HPV-positive (HPV+) HNSCCs have favourable prognoses and good treatment response rates, continuous exposure to tobacco smoke worsens the burden of mutations and leads to poorer outcomes. TP53 and other cell cycle and DNA repair key regulators are among the most frequent genetic alterations found in HNSCC [Chow L.A, 2020]. Furthermore, the HPV oncoproteins E6 and E7 lead to the degradation of p53 and the inhibition of the retinoblastoma-associated protein (RB1), respectively. Eventually, this may underlie an increase in the number of malignant cells, which are not eliminated due to p53 degradation and its associated decreased apoptotic signaling.

Radiation therapy (irradiation, IR), surgical resection, and systemic chemotherapy are prime modalities used in the management of patients with local or locoregionally-confined HNSCC. The application of IR is not limited only to early-stage disease but is also relevant for the treatment of advanced HNSCC stages as well as oligometastases [Bahig et al, 2022]. EGFR inhibitors, immunotherapeutic, and targeted agents are currently under broad clinical investigation in combination with IR [Anderson et al, 2021; Byeon et al, 2019]. The main concerns are the high toxicity of such a combined treatment regimen and the low efficacy gain compared to treatment-induced side effects. Radiosensitizing agents can potentially limit the toxicity of IR by decreasing its doses and volumes but maintaining, or potentially, also increasing its efficacy [Li Q et al, 2023].

Ionizing radiation (IR) generates multiple DNA double-strand breaks (DSBs), which can be physiologically repaired by either the more demanding, sister chromatid-depending and therefore error-free homologous recombination (HR) or by the error-prone non-homologous end joining (NHEJ) pathway [Vitor et al, 2020]. Peposertib (formerly M3814) is a potent and selective small-molecule inhibitor of the catalytic subunit of DNA-PK (DNA-PKcs), a key effector of NHEJ [van Bussel et al, 2021]. Up to date, multiple studies have shown that NHEJ inhibition increases the antitumor treatment efficacy of various DNA-damaging agents, including IR and topoisomerase II inhibitors in distinct cancer types [Wise et al, 2019; Haines et al, 2021]. For instance, peposertib demonstrated a strong radiosensitizing effect on human cancer cells of various origins in-vitro and in-vivo [Klein et al., 2017; Zenke et al., 2020] and an improved chemoradiation effect in some rectal cancer models [Smithson et al., 2022].

Taking into account the disadvantages of conventional therapies and the persisting limitations of advances in HNSCC management, we addressed here the need for a novel treatment approach for these cancers by inhibition of DNA-PKcs prior IR in the context of the cellular p53 status. We hypothesized that this combination may improve HNSCC treatment outcome due to a compromised repair of RT-induced DSBs, particularly in cancer cells with p53 and/or HPV alterations, which are compromised in the maintenance of the G1/S cell cycle checkpoint.

We show here that peposertib radiosensitizes HNSCC tumors regardless of their p53 and HPV status by significantly increasing IR-induced γH2AX and RAD51 foci formation and causing a delay in DNA repair. Furthermore, inhibition of NHEJ combined with IR results in different cell fates depending on p53 functionality. Whereas HPV+ and p53-mutated cells are eliminated by apoptosis due to a common alteration in p53-related pathways, p53 wild-type cells preferentially undergo senescence. Combining DNA-PKcs inhibition and irradiation leads to a better treatment response in HNSCC models with dysfunctional p53 both in-vitro and in-vivo, indicating, therefore, the beneficial outcome of this combination in HPV+ or p53-mutated cancers. These data provide the first observations that may suggest a precision medicine-based treatment for HNSCC, which takes into consideration the p53 and HPV status of the tumor.

## Materials and Methods

### Cell lines

UM-SCC-10B, -14A, -17B, -74A, and -104 squamous cell carcinoma cell lines (kindly provided by Prof. Thomas Carey (University of Michigan) were cultured in DMEM (Sigma, D6046) supplemented with 10% heat-inactivated Fetal Bovine Serum (FBS-hi; Sigma, Cat. no. F7524), 1% antibiotic-antimycotic (AA: penicillin 100 U/mL, streptomycin sulfate 100 U/mL, 412 amphotericin B 0.25 mg/mL; Gibco, Cat. no. 15240-062), and 1% non-essential amino acids (NEAA; Sigma, M7145). The UD-SCC-2 cell line was obtained from Prof. Dr. Henning Bier (University of München, Germany) and cultured in RPMI (Sigma, Cat. no. R8758) supplemented with 10% FBI hi, 1% AA and 2mM L-Glutamine (Sigma, Cat. no. 59202C). The cell line UPCI-SCC-154 was purchased from the German Collection of Microorganisms and Cell Cultures GmbH (DMSZ) and cultured in MEM (Sigma, Cat. no. 51412C) supplemented with 10% FBI hi, 1% AA, 1% NEAA, and 2 mM L-Glutamine. All cells were maintained at 37°C with a 5% CO_2_ supply. The UM-SCC-10B, -14A, -17B, and -74A cell lines have been authenticated by whole-exome sequencing and transcriptomic profiling [Nisa et al, 2018]. All cells used in the study were tested negative by PCR (IDEXX BioAnalytics) for Corynebacterium bovis, Corynebacterium sp. (HAC2), and Mycoplasma sp. No other cell line authentication was performed by the authors for this study.

### In-vitro treatments

The DNA-PKcs inhibitor (DNA-PKcsi) peposertib (the healthcare business of Merck KGaA, Darmstadt, Germany) was dissolved in DMSO (Sigma, Cat. no. D8418), aliquoted, and used at a final concentration of 300nM. Cells were treated 24 hours after seeding in all experiments. Cells were irradiated using a ^137^Cs source (Gammacell 40, MDS Nordion, Ottawa, ON, Canada) at a dose rate of 0.80 Gy/min. In all experiments and unless specified differently, cells were irradiated with a single dose of 4 Gy 30 min after incubation with DMSO or peposertib. Overall, four treatment groups were used: control (DMSO), peposertib (300nM, 30min incubation prior IR), 4Gy irradiation (IR), and peposertib (300nM, 30min incubation prior IR) in combination with 4Gy irradiation (peposertib+IR).

### Protein extraction and western blotting

Cells were lysed 1h after treatment in urea buffer (20mmol/L HEPES, pH 8.0; 9.0 mol/L urea; 1mmol/L sodium orthovanadate; 2.5mmol/L sodium pyrophosphate, 1mmmol/L β-glycerol-phosphate) and sonicated (Bandelin Sonopuls, HD2070). Total protein concentration was determined with the Bio-Rad protein quantification reagent (Bio-Rad Laboratories, Inc.). Equal amounts of protein (50 µg) were separated by SDS-PAGE and transferred to PVDF membranes. The membranes were blocked in 5% BSA (Roche, 10 735 086 001) in TBS-T before overnight incubation at 4°C with primary antibodies: pDNA-PKcs (Abcam, Cat. No. 124918), p53 (Cell Signaling Technology (CST), Cat. No. 9282), p-p53 (CST, Cat. No. 9284), p21 (Millipore, Cat. No. 05-655), cleaved Lamin A (CST, Cat.No 2035), β-Actin (Millipore Cat.No. MAB1501). Immunodetection was performed on an Odyssey imager (Li-Cor Biosciences) following incubation with secondary antibodies (IRDye 680RD goat anti-mouse, Li-Cor, Cat. no. 925-68070, and IRDye 800CW goat anti-rabbit, Li-Cor, Cat. no. 925-32211).

### Cell cycle assay

Cells were fixed 7, 24, 36, and 48 hours after the corresponding treatments in 70% EtOH, washed in PBS, and resuspended in 200µL propidium iodide (PI) solution (50µg PI + 40µg RNAse A/mL). PI incorporation was measured by flow cytometry (BD^TM^ LSR II, BD Biosciences) and analyzed using the FlowJo software (FlowJo, Ashland, OR).

### CFSE generation tracking

Generation tracking has been performed according to the manufacturer’s instructions (CellTrace CFSE, Invitrogen). Before plating, cells were stained with carboxyfluorescein succinimidyl ester (CFSE), a fluorescent dye, which covalently binds proteins and is equally distributed to the daughter cells upon division. Five days after treatment, the assay was terminated, and the intensity of the signal was measured using flow cytometry. On the day of treatment, an untreated sample was collected to serve as a basal intensity signal. Signal intensity was measured by flow cytometry using LSR II (BD Biosciences), and plotted with FlowJo (FlowJo, LLC).

### Cell proliferation assay

Cells were seeded in 24-well plates, treated with either DMSO or peposertib, exposed to different doses of IR (0, 2, and 4Gy), and then allowed to proliferate for 9 days. At the end of the experiment, the cells were fixed and stained with 2% crystal violet (SIGMA, C0775) dissolved in acetic acid:methanol (2:1). Cell density was calculated by determination of the covered area using ImageJ (imagej.nih.gov/ij/) and normalized to the corresponding non-irradiated control.

### Clonogenic assay

Single cells were seeded in triplicates in 6-well plates, treated with either DMSO or peposertib, and exposed to different doses of IR (0, 2, and 4Gy). Fourteen days after plating, cells were fixed and stained with 2% crystal violet (SIGMA, C0775) dissolved in acetic acid:methanol (2:1). Clonogenic growth was determined using Colcount, Charm Enhanced Algorithmus (Oxford Optronics, UK). The clonogenic fraction of irradiated cells was normalized to the corresponding non-irradiated control.

### XTT assay

Cells were plated into a 96-well plate in 50μL of the corresponding medium (six wells per treatment condition). After 24h, DMSO or peposertib solutions were added to reach the volume of 150μL, and the plate was irradiated (4Gy). 96 hours later, cellular proliferation was measured using the commercially available kit and the recommended protocol (Roche, Cat. no. 11465015001). Cells were incubated with the XTT reagent at 37°C, 5% CO_2_ for 2 hours. During the incubation, the yellow tetrazolium salt XTT is cleaved to formazan, the amount of which is correlated to the number of metabolically active cells. Quantification was performed by measuring the absorbance at 490 nm (reference wavelength: 655 nm) using a Tecan Plate reader and relative metabolic activity was normalized to the corresponding nonirradiated control.

### Cell viability and toxicity assay

Cell death and viability were assessed by the Live/Dead Assay Kit (Molecular Probes, Cat. no. L23105) according to the manufacturer’s instructions. Briefly, cells were allowed to grow for 96 hours after treatment and then stained with green fluorescent calcein-AM (final concentration (f.c.) 1.6µM) and red fluorescent ethidium homodimer-1 (f.c. 4µM) to detect viable and dead cells, respectively. Cells were imaged with a fluorescence microscope (Leica 4000) at 4x magnification and quantified with ImageJ.

### Caspase-3 activity assay

Cells were lysed in assay buffer (100 mM HEPES, pH 7.5, 1% sucrose, 0.1% CHAPS) 72h post-treatment, and cell protein content was determined with the Bio-Rad protein quantification reagent (Bio-Rad Laboratories, Inc.). Caspase-3 activity was assessed in duplicates via a fluorogenic assay using the Ac-DEVD-AMC-specific caspase-3 substrate (Calbiochem, La Jolla, CA). After substrate addition, fluorescence was measured with a TECAN Infinite200 pro plate reader (Männedorf, Switzerland), and the signal of caspase-3 activity was normalized to the samples’ protein content and the corresponding nonirradiated control.

### Senescence-associated β-galactosidase assay

Cells seeded and treated in 8-chamber slides (Falcon, Cat. no. 354118) were stained 9 days after treatment. Reagents and staining for senescence-associated β-galactosidase were implemented as described previously [Debacq-Chainiaux, 2009]. In short, the cells were fixed for 7 minutes in 2% formaldehyde/0.2% glutaraldehyde solution at room temperature and further incubated with X-Gal (Sigma, Cat. no. B4252) solution for one hour at 37°C. All washes were done with PBS. Images were obtained with an inverted fluorescent microscope (Leica 4000) with 63x immersion objective magnification.

### γH2AX and RAD51 foci immunostaining

Cells were seeded in 8-chamber microscopy slides and fixed in 4% formaldehyde (SIGMA, 158127) at 3, 24, and 48 hours after the corresponding treatment. After fixation, the cells were permeabilized with 0.1% Triton x-100 (Fluka, 93426) and blocked with the blocking solution containing 3% Goat serum (Dako, Cat. no. X0907), 0,1% sodium azide (the healthcare business of Merck KGaA, Darmstadt, Germany, Cat. no. 6688), 0,5% casein (Sigma, Cat. no. C-8654), 0,025% Tween (Sigma, Cat. no. P2287) dissolved in TBS pH 7,5. Washes were performed with TBS and TBST. The cells were incubated overnight with primary antibodies targeting γH2AX (CST, Cat. no. 2577S), and RAD51 (GeneTex, Cat. no. GTX70230) foci. Slides were mounted with Vectashield antifade mounting medium (Vector Laboratories, Cat. no. H-1000) and prepared for imaging with an inverted fluorescent microscope (Leica 4000) with 63x immersion objective magnification after 1h incubation with secondary antibodies (goat anti-mouse IgG 546 (Molecular Probes, Cat. no. A11030) and goat anti-rabbit IgG 488 (Molecular Probes, Cat. no. A11008)) and counterstain with DAPI (Sigma, Cat. no. D-9542). Quantification of foci per nuclei was performed with a modified pipeline used for foci count on CellProfiler (www.cellprofiler.org) software.

### Mouse xenograft model

Animal experiments were performed in the Central Animal Facility of the University of Bern following the general guidelines and regulations. All experiments were approved by the local experimental animal committee of the Canton of Bern according to Swiss laws for animal protection (animal license BE128-20). Mouse xenograft models employing UM-SCC-74A (p53 wild-type) and UD-SCC-2 (HPV+) cell lines were established. Two million cells diluted in 50ul solution of Matrigel® Basement Membrane Matrix (Corning, Cat. no. 356237) and Dulbecco’s phosphate-buffered saline (1:1) were injected subcutaneously to the right posterior flank region of five-week-old male and female NMRI-nu mice (Janvier Labs, France). Tumor formation was monitored twice a week by palpation and calliper measurement. Tumor volumes were calculated employing the following formula: tumor volume = (A^2^*B)/2, where A is the diameter of the largest measurement and B is the diameter of the smallest measurement. Once the tumors reached a volume between 100 to 300 mm^3^, mice were randomized to four different treatment groups (vehicle, peposertib, IR, and peposertib+IR, n=7 in each group). Since the first day of the treatment, the regular follow-up including weight and tumor measurement was maintained for 45 days or until the euthanasia criteria (e.g., tumor size is above 1000mm^3^) were met. At the end of the follow-up, the mice were euthanized with pentobarbital (150 mg/kg, Esconarkon, Streuli Pharma AG), and tumor tissues were collected. Each tumor was half dissected and fixed in either 10% buffered formalin (Simport, Cat. no. M960-40FMA) or freshly cryopreserved in the isopentane (Sigma. Cat. no. M32631) to proceed with immunostaining.

### In-vivo models and interventions

Peposertib powder was suspended in a vehicle solution containing Methocel 0,5% (Sigma, Cat. no. 94378), Tween-20 0,25% (Sigma, Cat. no. P2287), and Na-Citrate buffer 300mM (pH=2,5). The vehicle solution alone was administered orally via a gavage needle with a volume of 200ml for five consecutive days. Peposertib dissolved in vehicle solution was delivered in the same way with a single dose of 100mg/kg (cumulative dose was 500mg/kg). Two Gy of irradiation per fraction was delivered precisely to the tumor site for five consecutive days (cumulative dose of 10 Gy) by the small animal radiation therapy system (SmART; Precision X-Ray Inc.). While combining the two perturbations, the drug was given 15 minutes before radiation therapy. Mice were anesthetized in the induction chamber with a mix of 5% isoflurane and 100% oxygen (1L/min) and then moved to the animal stage of the SmART machine where the anesthetic flow was maintained via face mask throughout the whole procedure. For each mouse, the single irradiation dose (2Gy) was calculated by Monte Carlo dose calculation (SmART-ATP) on the first day of treatment based on the acquired high-resolution CT scan (PilotXRAD software) of the subcutaneous tumor. The delivery was performed using a SmART tube (collimator field size – 10mm*10mm, filter: 5 mm Cu) from two beaming angles (180°). On treatment days 2-5, the radiation was delivered with the same plan using CT-guided tumor targeting.

### Tissue staining

Formalin-fixed paraffin-embedded (FFPE) sections of 5 µm thickness and fresh cryosections of 10 µm were prepared from the xenograft tumors. The first half of the FFPE sections were deparaffinized, immersed in a citrate buffer for 30min at 95-100°C, incubated with peroxidase blocking solution 3% H_2_O_2_ (Millipore Sigma, 107209), and blocked with 1% Normal Goat Serum (DAKO, Cat. no. X0907). Washes were performed with TBS and TBST. The slices were incubated overnight at +4°C with a primary antibody targeting Ki67 (CST, Cat. no. 9027S). After 1h incubation with Goat biotinylated anti-rabbit IgG (Vector, Cat. no. PK-6101) at room temperature, the sections were incubated with ABC reagent solution (Vector, Cat. no. PK-6101) for 30min and with DAB solution (Vector, Cat. no. SK-4100) for 4 minutes. Afterwards, the slices were counterstained with hematoxylin, dehydrated, and cover-slipped with Eukitt mounting medium (Sigma, Cat. no. 03989). The second half of the slices were stained for apoptosis assessment according to the DeadEnd™ Fluorometric TUNEL System (Promega, Cat. no. G3250) protocol. In short, the sections were deparaffinized, fixed in 4% formaldehyde, permeabilized with 20µg/ml Proteinase K solution, equilibrated, and incubated with TdT for 60 minutes at 37°C in a humidified chamber. The reaction was stopped by immersing the slides in 2X SSC, counterstained with DAPI, and mounted with Vectashield antifade mounting medium. Washings were performed with 0.85% NaCl and PBS. The fresh cryopreserved sections were spared for beta-galactosidase senescence assay. Tumor slices were fixed with 2% Glutaraldehyde (Sigma, Cat. no. G6257) for 15 minutes at room temperature. The sections were incubated with freshly prepared β-gal solution (Sigma, Cat. no. B4252) overnight at 37°C. The slides were counterstained with hematoxylin and coverslipped with Vectashield antifade mounting medium. The washes were done with PBS. All slides were light-protected and stored at +4°C before imaging with a Slide scanner (Pannoramic 250 Flash II, 3DHistech) and the quantification was performed with the Fiji package (Image J).

### Statistical analysis

Statistical analysis and the graphical presentation of the data were performed using GraphPad PRISM (GraphPad Software Inc, version 9.5.0). Some of the elements of the figures were created with BioRender.com. One-way or two-way ANOVA with post hoc tests have been performed. Log-rank (Mantel-Cox) test has been performed for survival analysis. All experiments were replicated independently three times unless indicated differently. The data in the text is presented as relative averages or fold-induction ± SD, as indicated. The bars in the graphs are represented as means ± S.E.M. Differences between the groups were considered significant with p values <0.05 (ns – non-significant, * p<0.05; ** p<0.01; *** p<0.001, **** p<0.0001).

## Results

### DNA-PKcs phosphorylation in head and neck cancer cells following IR occurs independently of their HPV and p53 status and can be inhibited by peposertib

Radiotherapy (RT) with or without concomitant chemotherapy is a standard therapeutic option for most of head and neck cancers. As these treatments cause early and late toxicities and their outcomes vary substantially depending on factors such as tumor subtype, localization, or molecular composition, novel therapeutic options including rational combination approaches are direly needed. Here we aimed to study the efficacy of IR in combination with the DNA-PKcs inhibitor peposertib in HNSCC cell lines that differ in their HPV and p53 status. Whereas TP53 mutations are widely prevalent in HPV-negative (HPV-) head and neck cancers [de Bakker et al., 2022], in HPV+ tumors, the E6 oncoprotein of HPV leads to the degradation of p53 via an E3-ubiquitin ligase and hence prevents the detection and elimination of abnormal and potentially cancerous cells by p53 [Wiest et al., 2002]. We have assessed basal and RT-induced DNA-PKcs Ser2056 and p53 Ser15 phosphorylation as well as total protein expression of p53 and p21 in a panel of 7 HNSCC cell lines including HPV-models with either functional or mutated p53 as well as HPV-positive ones (Fig. 1A). We detected basal p53 and p21 protein levels in the two p53 wild-type cell lines (UM-SCC-17B and UM-SCC-74A) and an increase in both total and Ser15-phosphorylated levels of p53 after IR. Cell lines bearing either a missense (UM-SCC-10B [Sano et al., 2011] or nonsense (and UM-SCC-14A [Ludwig et al., 2019]) p53 mutation also express the p21 protein and comparably high basal levels of total p53 but no basal or radiation treatment-induced p53 Ser15 phosphorylation can be detected (Fig. 1A). Lastly, HPV+ cell lines (UD-SCC-2, UM-SCC-104, and UPCI-SCC-154) express neither total nor phosphorylated p53 and negligible p21 levels (Fig. 1A).

**Figure 1.**
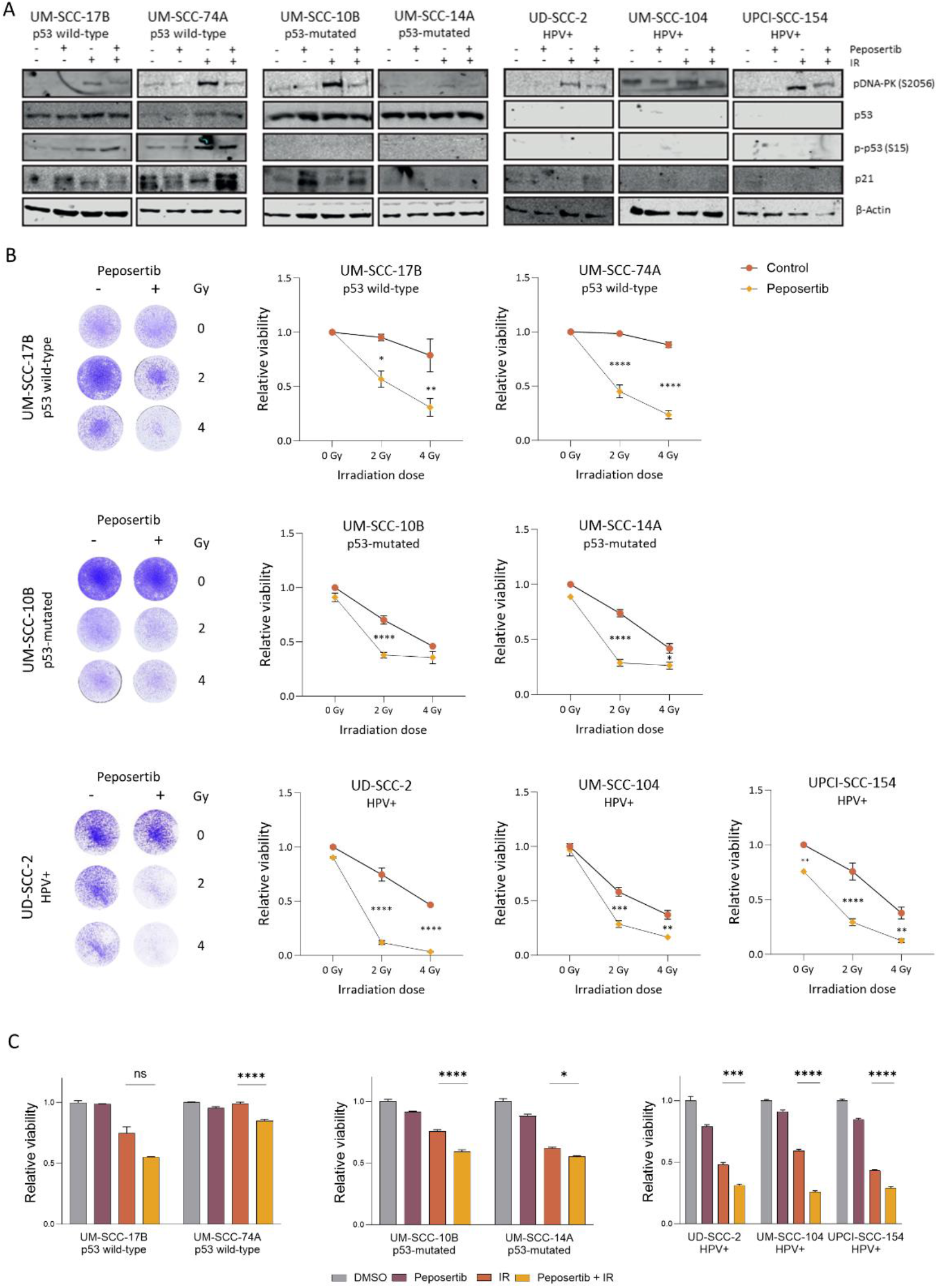
DNA-PKcs can be targeted in HNSCC, and its inhibition prior IR causes cell cycle redistribution in p53-mutated and HPV+ HNSCC models. **(A)** Phosphorylation levels of DNA-PKcs (S2065) and p53 (S15) and total protein levels of p53 and p21 in whole-cell lysates of untreated p53 wild-type (UM-SCC-17B, UM-SCC-74A), p53-mutated (UM-SCC-10B, UM-SCC-14A) and HPV+ (UD-SCC-2, UM-SCC-104, and UPCI-SCC-154) cell lines as well as in cells treated with peposertib (300nM, 30 minutes incubation), IR (4 Gy, lysis 1h post treatment) or their combination. ß-Actin was used as a loading control. **(B)** Proliferation of HNSCC cell lines assessed 5 days after treatment (control, peposertib, IR or their combination) by a crystal violet assay *(left* -representative images, *right* - quantification (data is represented as *mean±SEM;* two-way ANOVA with post-hoc multiple comparisons: *\* p<0.05, ** p<0.01, *** p<0.001, **** p<0.0001;*). **(C)** Viability of distinct HNSCC cell lines upon treatments by peposertib, IR and their combination assessed by the XTT assay (Data is represented as *mean±SEM;* one-way ANOVA with post-hoc multiple comparisons: *ns – nonsignificant, * p<0.05, ** p<0.01, *** p<0.001, **** p<0.0001*. Significance shown only for comparisons between peposertib+IR and IR alone.)

Importantly, DNA-PKcs Ser2056 phosphorylation was induced after irradiation in all the studied cell lines, thus indicating the activation of repair of DNA DSBs via the NHEJ pathway (Fig. 1A). At the same time, when the selective DNA-PKcs inhibitor peposertib was administered 30 minutes before RT, the DNA-PKcs Ser2056 phosphorylation levels were considerably lower across the entire cell line panel as compared to IR treatment alone. We also observed a DNA-PKcs inhibition-dependent and p53-independent increase in basal and irradiated p21 levels in the p53-mutated cell line UM-SCC-10B and both p53 wild-type cell lines (Fig. 1A).

### The efficacy of peposertib-mediated HNSCC radiosensitization differs between p53 wild-type, p53-mutated, and HPV-positive cell lines

To investigate how DNA-PKcs inhibition before IR impacts viability and survival of HNSCC cancer cell lines with distinct HPV and p53 status, we performed crystal violet staining (Fig. 1B and Suppl. Fig. 1) and colony forming assays (Suppl. Fig. 2C), respectively. We did not observe a significant decrease in viability upon peposertib treatment alone compared to the control condition in any of the cell lines except for the HPV+ UPCI-SCC-154 cells where the viability was reduced by 24.3%. On the contrary, IR alone decreased the viability of all cell models included albeit to a different extent. A single dose of 2 or 4 Gy reduced the viability of HPV-positive and p53-mutated by 29.5 and 58.2% whereas of the p53 wild-type cells only by 3.1 and 16.6%, respectively. Importantly, pretreatment by the DNA-PKcs inhibitor peposertib 30 minutes before 2Gy of IR compared to IR alone decreased the viability significantly in all cell lines whereas the combination treatment using 4 Gy had a similar effect solely in p53 wild-type and HPV+ HNSCC cell lines, most probably due to the already high radiation sensitivity of p53-mutated cell lines at this dose.

In line with these findings, the administration of peposertib alone does not impair the ability to form colonies in any of the cell lines, whereas already 2 Gy of IR provokes a decreased survival rate. When the cells are treated with both peposertib and 2 Gy of IR, clonogenic survival of most of them is further impaired (Suppl. Fig. 2D).

Besides, we screened our panel of cell lines for metabolic activity using an XTT assay (Fig. 1C). Similarly to the data resulting from the crystal violet and colony-forming assays, peposertib alone does not significantly impact the metabolic activity of any of the studied cell lines. However, its combination with IR significantly radiosensitizes all the cells apart of the p53 wild-type-expressing UM-SCC-17B cell line, where we detected a substantial but not significant reduction in metabolic activity. In addition, HPV+ cell lines seem to be more sensitive to both IR and combined peposertib+IR treatments compared to p53-mutated HNSCC cells and cells with functional p53. Noteworthy, we could observe a higher fraction of dead cells after IR and a combination of IR and peposertib compared to the control in all cell lines (Suppl. Fig. 3).

### Combination of DNA-PKcs inhibition and IR arrests p53-depleted cell lines in the G2 phase of the cell cycle

As HPV inactivates p53 to reinitiate cell division after terminal differentiation, we aimed to investigate whether HNSCC cell lines with distinct p53 and HPV status display different cell cycle distribution patterns upon DNA-PKcs inhibition before IR. Our results reveal that HPV-positive (UD-SCC-2 – 73.1%±10.6 (vs 12.8%±6.9, p<0.0001), UM-SCC-104 – 46.2%±11.1 (vs 19.5%±4.3, p=0.0003) and UPCI-SCC-154 – 49.3%±12.1 (vs 18.8%±3.1, p=0.0002)) and p53-mutated cell lines (UM-SCC-10B – 36.8%±7.0 (vs 15.6%±3.7, p=0.0020) and UM-SCC-14A – 64.8%±2.9 (vs 13.3%±0.7, p<0.0001)) treated with peposertib before IR strongly accumulate in the G2 phase of the cell cycle over a time course of 48 hours compared to their respective untreated controls (Fig 2A). Notably, G2 accumulation of HPV-positive and p53-mutated cells exposed to IR alone peaks around 24 to 36 hours before being released and steadily decreases to basal levels (Suppl. Fig. 2A-C). On the contrary, IR in combination with the DNA-PKcs inhibitor peposertib does not induce any G2 arrest in p53 wild-type cell lines at 48h after treatment (UM-SCC-17B – 26.6%±14.7 (vs 15.6%±3.7, p=0.0044) and UM-SCC-74A – 14.2%±6.3 (vs 16.05%±0.9, p=0.6746)) or at earlier time points.

**Figure 2.**
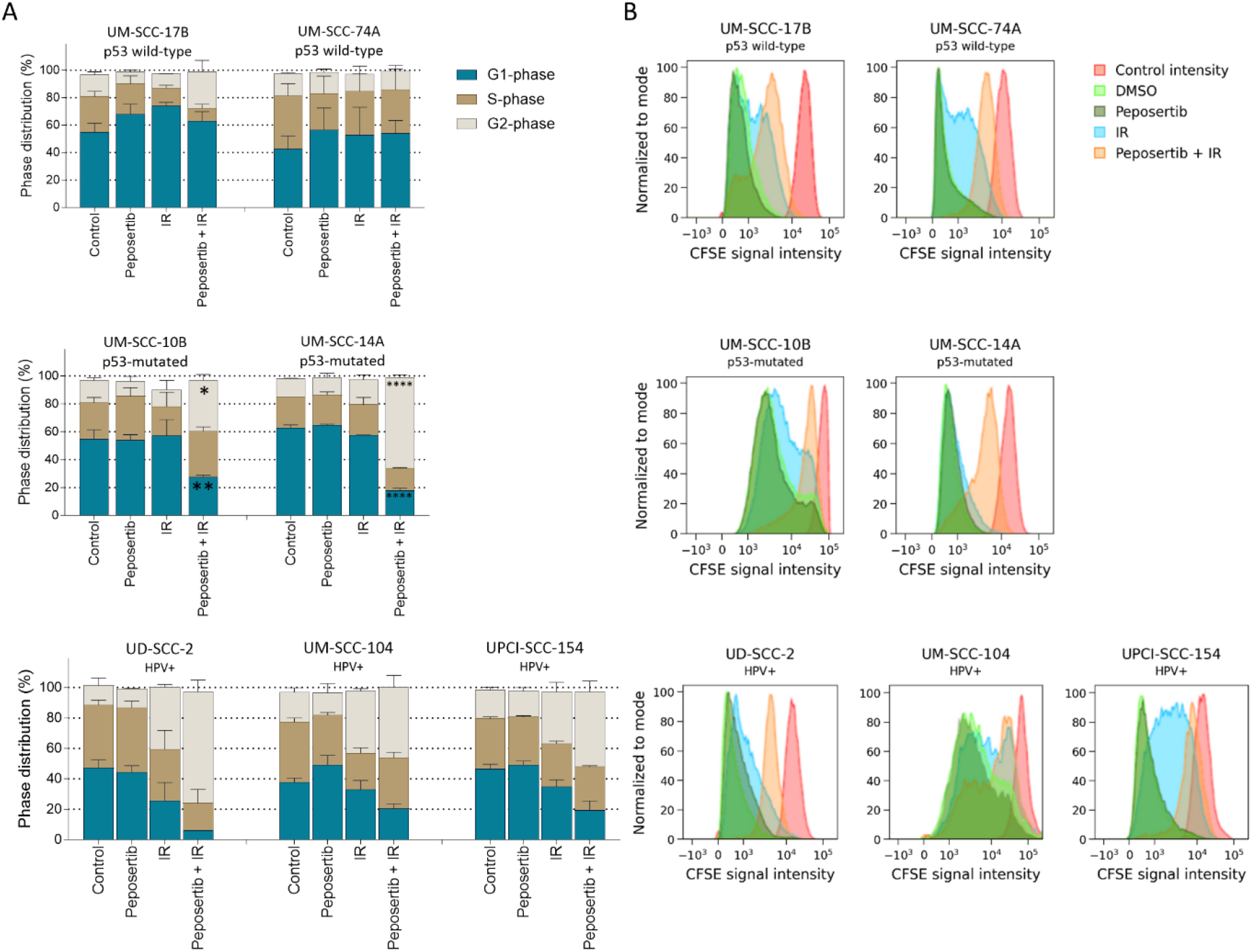
Combination of DNA-PKcs inhibition and IR arrests p53-depleted cell lines in the G2 phase of the cell cycle. **(A)** Cell cycle distribution in untreated cells and 48h upon treatment with peposertib, IR or their combination (data is represented as mean±SEM, significance was assessed by two-way ANOVA with post hoc multiple comparisons and is shown only for comparisons between peposertib+IR and control within the same phase: p<0.05, ** p<0.01, *** p<0.001, **** p<0.0001 **(B)** Carboxyfluorescein succinimidyl ester (CFSE) assay performed in HNSCC cell lines 5 days after treatment by DNA-PKcs inhibition, IR or their combination (*red peak* - control intensity, *light green* - control cells treated by DMSO, *dark green* - peposertib-treated cells, *blue* - IR-treated cells, *orange* - peposertib + IR treatment).

We also performed a generation tracking experiment to clarify whether the cell cycle arrest of HPV+ and p53-mutated cells is terminal and if p53 wild-type cell lines are arrested in the cell phase they reside in (Fig. 2B). In this assay, a stronger curve shift from the basal intensity signal suggests higher proliferation due to dye loss in dividing daughter cells. As seen in Fig. 2B, all HNSCC cell lines treated with peposertib alone proliferate at the same rate as untreated controls and they partially or completely recover from IR-inflicted damage, restore their proliferative capacity, and re-enter the cell cycle. On the contrary, almost a complete arrest of proliferation in all cell lines after combining the drug with IR can be observed. This suggests that although a particular genetic background promotes diverse cell phase distribution patterns, cell division after a combination of IR and DNA-PKcs inhibition is impaired independently of p53 status.

### Peposertib- and IR-induced abrogation of proliferation promotes cell fates depending on the genetic background

Sun et al. reported previously that p53 functionality impacts the mode of cell death upon DNA-PKcs inhibition in combination with IR in cancer cells of various origins [Sun et al, 2019]. Hence, we next aimed to determine whether the distinct patterns of viability and cell cycle distribution that we observed in our panel of HPV+ and HPV-HNSCC models lead to different cellular fates. As p53 triggers apoptosis and controls the balance between activation of pro-senescent and pro-apoptotic pathways [Mijit et al, 2020], we assessed the induction of apoptosis (Fig. 3A and B) and senescence (Fig. 3C) following the various treatments in the cell line panel. Our data indicate that only cell lines lacking functional p53 (e.g., HPV+ and p53-mutated cells) undergo apoptosis after IR and/or after the combination of IR and DNA-PKcs inhibition as seen from increases in cleaved lamin A levels (Fig. 3A) and caspase-3 activity (Fig. 3B). On the contrary, no lamin A cleavage or caspase-3 activity can be observed following these treatments in neither of the two p53 wild-type cell line, indicating that these cells completely circumvent activation of the caspase cascade. Instead, a significant increase in the number of senescent cells can be visualized by the cytochemical β-galactosidase assay solely in the two p53 wild-type cells UM-SCC-17B and UM-SCC-74A upon IR and this increase is yet significantly substantiated when peposertib treatment is used prior IR (Fig. 3C).

**Figure 3.**
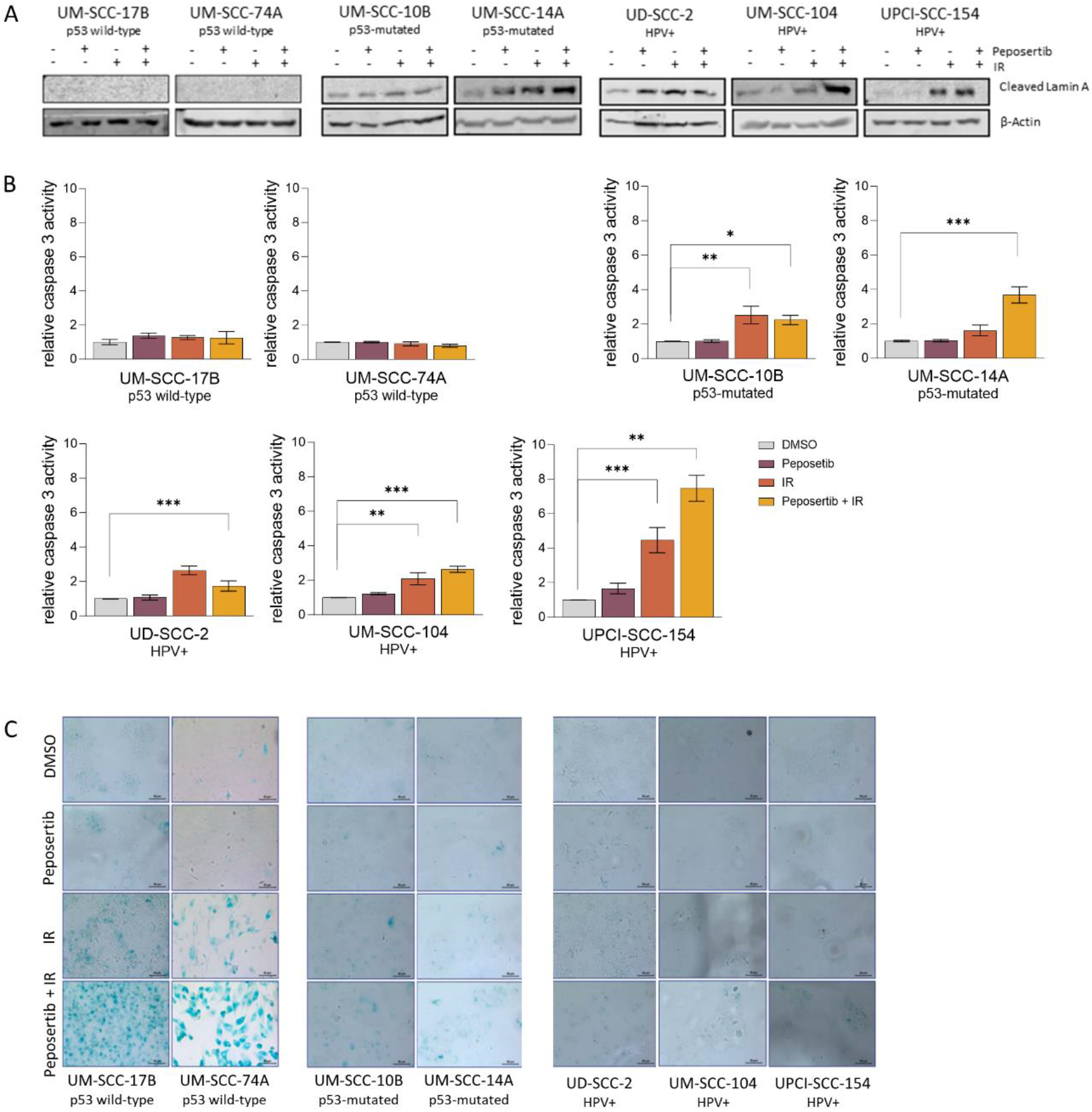
IR alone and in combination with peposertib induces apoptosis and senescence in cells with intact and compromised p53 signaling, respectively. **(A)** Cleaved Lamin A assessed in the HNSCC panel 1h post-treatment. ß-Actin was used as a loading control. **(B)** Caspase-3 activity assessed 72h post-treatment (Data is represented as mean±SEM; one-way ANOVA with post hoc multiple comparisons: *p<0.05, ** p<0.01, *** p<0.001, **** p<0.0001*)). **(C)** β-Galactosidase staining in HNSCC cell lines performed 9 days post-treatment. Representative images are shown.

### Peposertib administration before IR delays DNA repair in HNSCC cell lines regardless of their genetic background

To assess the level of DNA damage in HNSCC after various perturbations and to estimate the capacity of the cells to repair the damaged DNA via homologous recombination (HR) when NHEJ is blocked by peposertib treatment, we performed immunostaining targeting γH2AX and RAD51, respectively, 3h, 24h and 48h post-treatment (Fig. 4). We did not observe any significant differences in γH2AX or RAD51 foci formation between the cell line groups at 3h post-treatment: there was neither an increase in foci count upon peposertib distribution alone as compared to unperturbed cells nor any impact of the drug on the elevated γH2AX and RAD51 foci formation upon IR. However, a dramatic and highly significant increase in the number of γH2AX and RAD51 foci after combining DNA-PKcs inhibition and IR compared to the IR alone group can be observed almost uniformly across all the studied cell lines at later time points, especially 48h post-treatment. Altogether these data suggest that DNA-PKcs inhibition does not affect basal DNA damage levels in HNSCC in-vitro models but at the same time significantly increases IR-induced γH2AX foci formation as well as RAD51 activity, and this occurs independently of HPV or p53 status of the cells.

**Figure 4.**
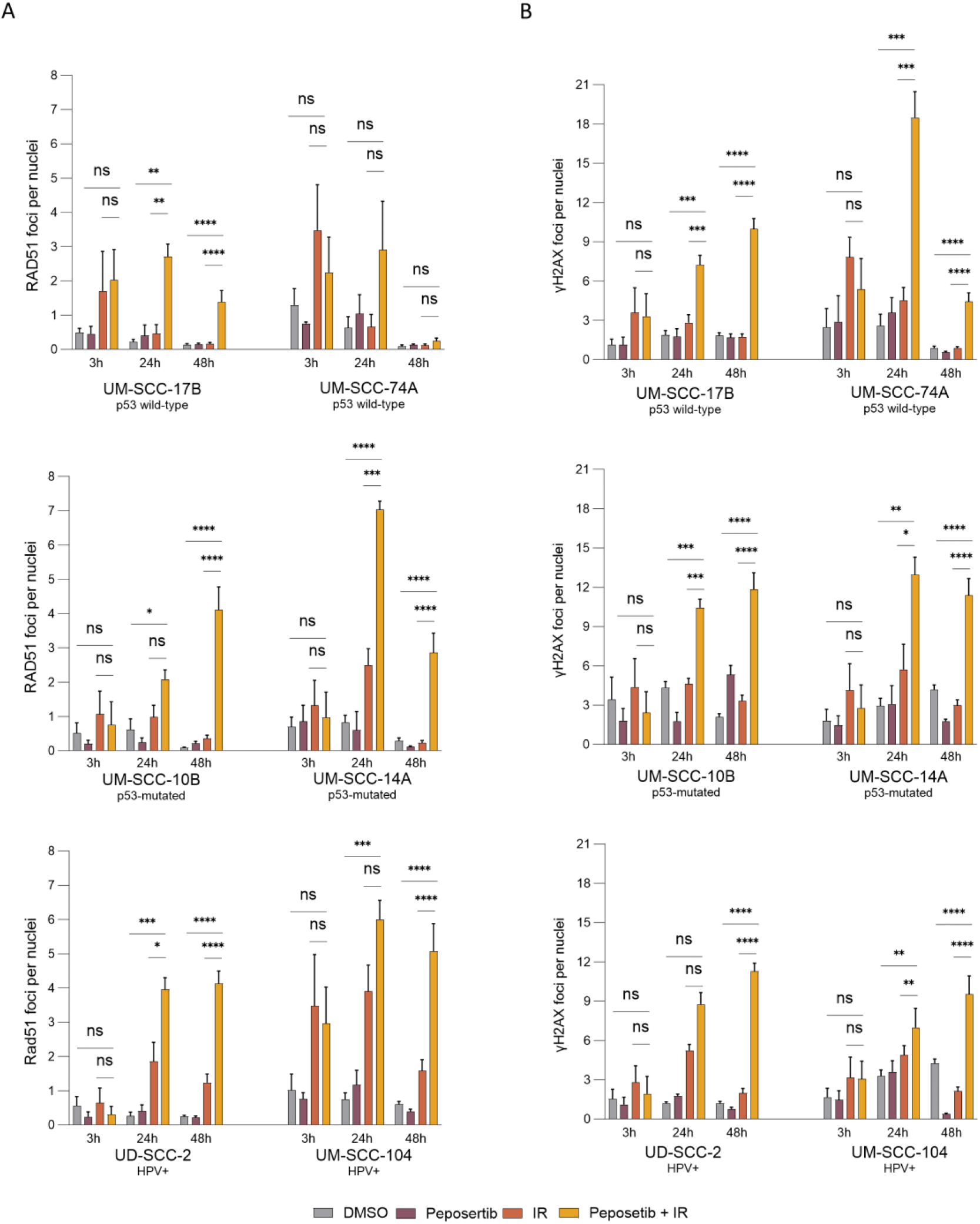
DNA-PKcs inhibition prior to IR delays DNA repair. **(A)**. Representative images of RAD51 and γH2AX foci in the nucleus. **(B)** RAD51 and **(C)** γH2AX foci quantified at 3h, 24h, and 48h after treatment. *Data has been analyzed with two-way ANOVA with post hoc multiple comparisons and is represented as mean±SEM; Significant marks stand for the comparison between peposertib+IR and control or IR alone: ns – nonsignificant, p<0.05, ** p<0.01, *** p<0.001, **** p<0.0001*

### HPV+ HNSCC xenografts respond significantly better to the combination of DNA-PKcs inhibition and IR compared to p53 wild-type tumors

To study the response of the HNSCC cell lines with distinct HPV backgrounds in-vivo, we have developed an HPV+ UD-SCC-2 as well as a p53 wild-type UM-SCC-74A subcutaneous xenograft mouse model (Fig. 5A). Once the tumors reached the size of 150-300mm^3^, mice were randomized to four different treatment groups (Suppl. Fig. 4A), which included fractionated IR regimen with a cumulative dose of 10Gy alone or in combination with peposertib administered orally before IR for five consecutive days. Importantly, the assessment of weight, overall mouse condition, and visual skin inspection did not indicate any toxicity arising from either peposertib, IR, or their combined treatment in any of the animals (Suppl. Fig. 4C). Although we did not find any significant differences in tumor growth between vehicle and peposertib-treated groups in both HPV+ and p53 wild-type xenografts (Fig. 5B and Suppl. Fig. 4B), IR treatment alone and DNA-PKcs inhibition in combination with IR significantly impacted tumor sizes in both animal models throughout the whole follow-up. Interestingly however, whereas DNA-PKcs inhibition in combination with IR (V = 232.1mm^3^±324.0) compared to IR alone (V = 1098.1mm^3^±400.0) caused a significant and sustained tumor growth suppression in the HPV+ UD-SCC-2 xenografts, the p53 wild-type UM-SCC-74A tumors, whose growth was initially strongly delayed by the combination treatment (V = 641.0mm^3^±576.1 and V(IR) = 956.9mm^3^±413.1), eventually relapsed throughout the follow-up. Consequently, we did not observe any significant differences in UM-SCC-74A tumor sizes following IR treatment in the presence or absence of DNA-PKcs inhibition on the day of experiment termination. The same pattern has been documented by survival analysis (Fig. 5C). These findings point towards a possibly high clinical benefit of DNA-PKcs inhibition in combination with IR specifically in HPV+ HNSCCs, thus reinforcing the conclusions of the in-vitro data.

**Figure 5.**
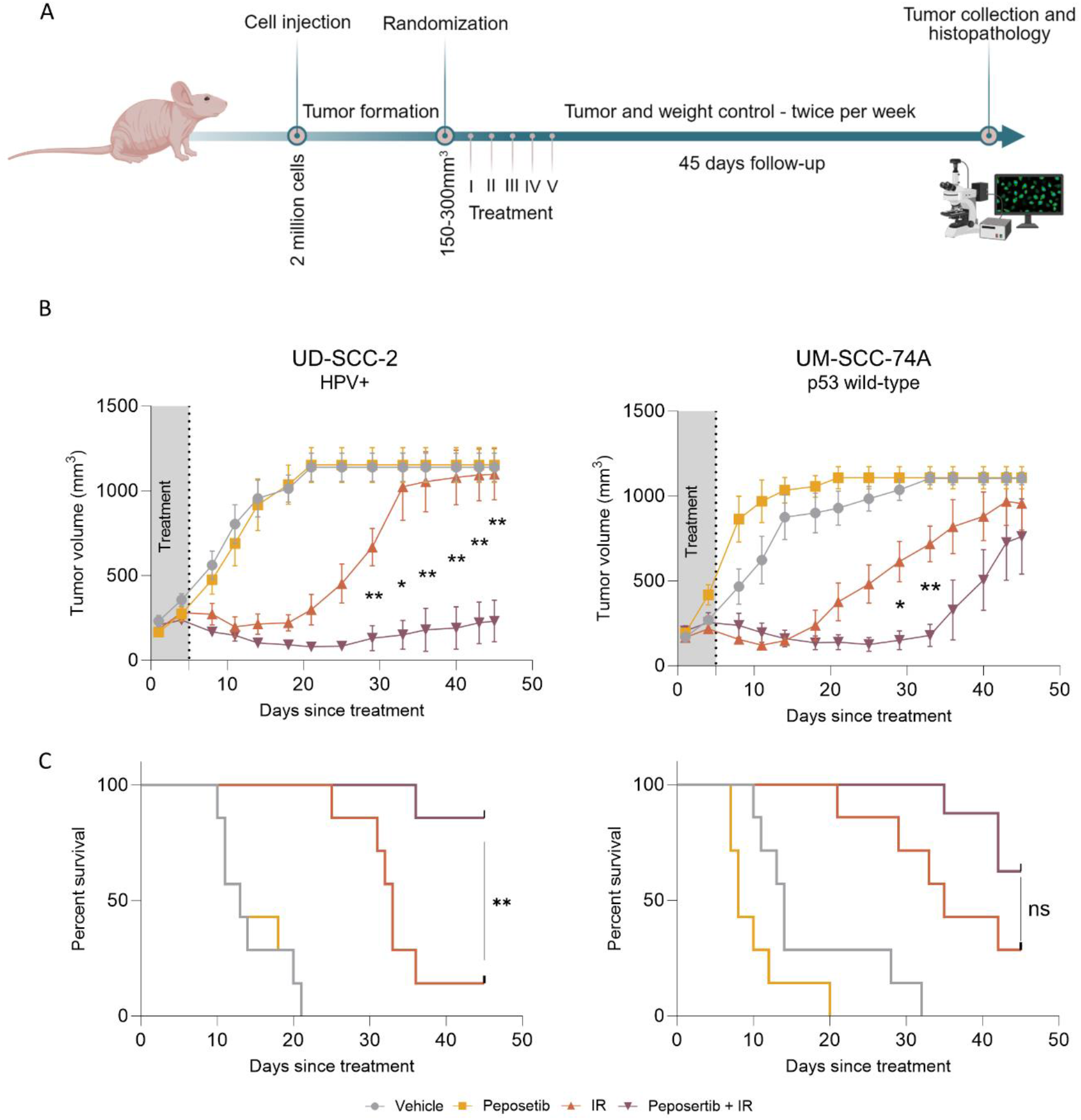
DNA-PKcs inhibition prior to IR effectively suppresses tumor growth in HPV+ xenograft tumors. **(A)** Timeline of the in-vivo experimental setup. **(B)** Tumor volume throughout the follow-up (n=7, data is represented as mean±SEM; *two-way ANOVA with post hoc multiple comparisons*). **(C)** Survival plot based on the tumor harvest time point (Log-rank (Mantel-Cox) test has been performed for the survival analysis, X-axis – time, y-axis – 1 = event (tumor harvest), 0 = censored subject (day 45 post-treatment)). *Significant marks stand for the comparison between peposertib+IR and IR at the given time point or the end of the experiment: ns – nonsignificant, p<0.05, ** p<0.01, *** p<0.001, **** p<0.0001*

### HPV+ xenografts are cleared by apoptosis whereas peposertib-treated p53 wild-type tumors contain senescent cells

To determine if the distinct cell fates observed following the combined DNA-PKcs inhibition and IR treatment in-vitro are possibly recapitulated also in the in-vivo models, we assessed the presence of proliferating, apoptotic and senescent cells in tumor tissues collected at the end of the experiment by performing immunostaining targeting Ki67 (Fig. 6A), TUNEL staining for nuclear DNA fragmentation (Fig. 6B), and β-gal assay (Fig. 6C), respectively. We did not find any major differences in Ki67 expression and the number of fragmented DNA between the treatment groups within the same tumor type. Nevertheless, and in accordance with the in-vitro data, we observed a significantly higher basal level of apoptotic cells in the HPV+ tumors compared to the p53 wild-type xenografts (33.7% and 3.3%, respectively). Interestingly, the p53 wild-type UM-SCC-74A tissues stained positively for β-Gal when exposed to either peposertib alone (82.5% β-gal-positive cells (tumors collected on average on day 10.7 post-treatment start)) or in combination with IR (9.7% β-gal-positive cells (tumors collected on average on day 43.4 post-treatment start)). Whereas we could demonstrate the presence of β-gal-positive cells upon the combined treatment also in-vitro, the high number of senescent cells found in UM-SCC-74A tumor tissues following DNA-PKcs inhibition alone may indicate that peposertib treatment affects cancer cell fitness in the long-term.

**Figure 6.**
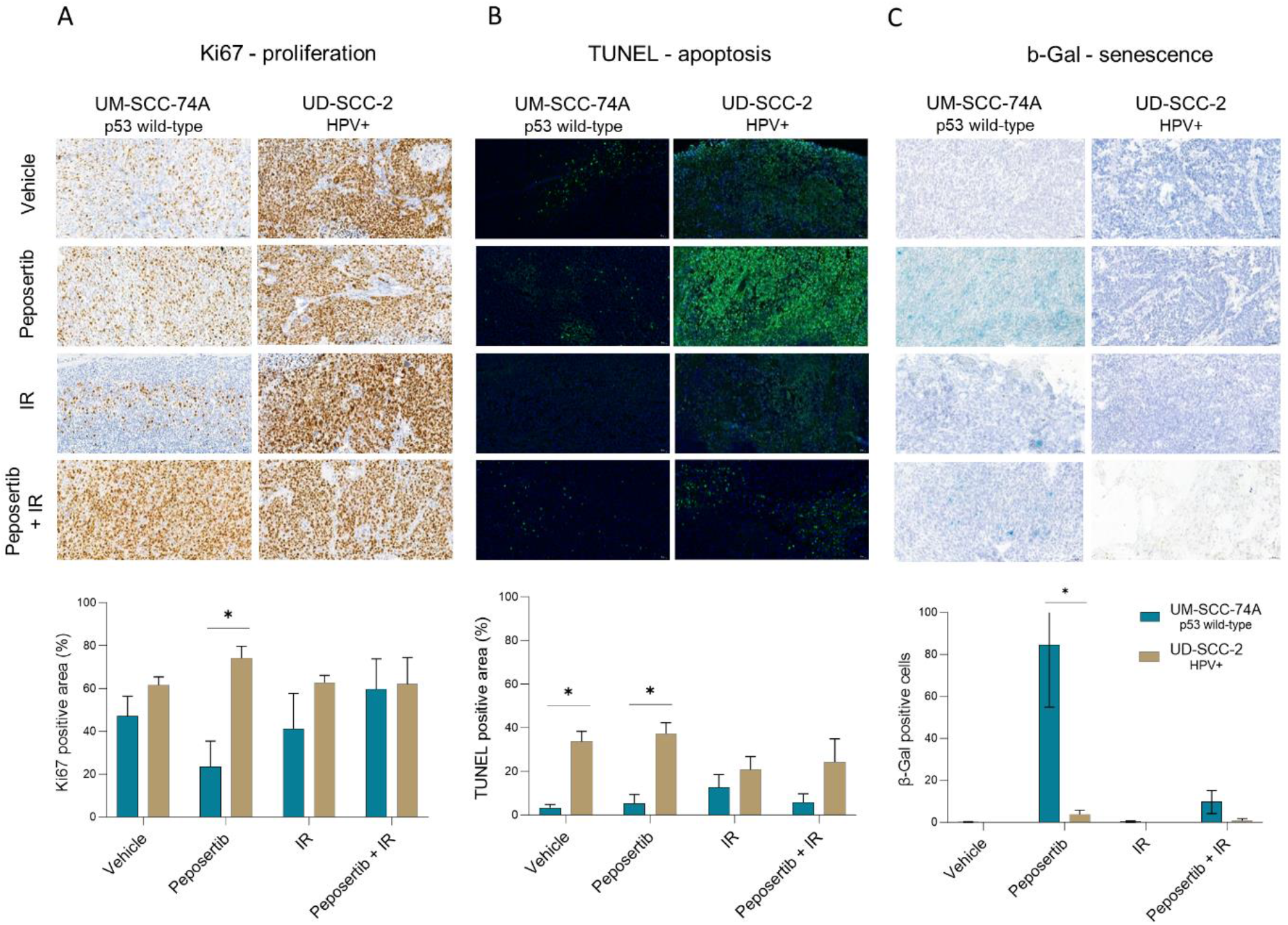
HPV+ xenograft tissues exhibit a higher basal count of apoptotic cells and a lower number of peposertib-treated senescent cells as compared to HPV-tissues. Immunostaining of **(A)** Ki67 (marker of proliferation), **(B)** nuclear DNA fragmentation (TUNEL staining), and **(C)** β-gal (marker of senescence). *Data has been analyzed with two-way ANOVA with post hoc multiple comparisons and is represented as mean±SEM; ns – nonsignificant, p<0.05, ** p<0.01, *** p<0.001, **** p<0.0001*

## Discussion

Although cure rates of HNSCC patients diagnosed at an early stage are as high as 80% [Johnson et al., 2020], management of cancer patients with locally advanced tumors is very challenging, and novel treatment approaches that would reduce the risk of recurrence without increasing toxicities are needed [Li et al., 2023]. Identification of molecular markers that shape tumor response to therapeutic interventions and could be used for patient stratification, may significantly improve treatment outcome. In this study, we assessed the effectiveness of NHEJ blocking in combination with IR in HNSCC models. As TP53 mutations and HPV positivity are two molecular characteristics commonly found in HNSCC tumors (60-80% and 84%, respectively) [Leemans et al, 2018], we also investigated how HPV and p53 status affect the outcome of this intervention.

Our data confirm that DNA-PKcs is a valid target in irradiated HNSCC models as we observed an IR-induced increase in DNA-PKcs Ser2056 autophosphorylation, an established pharmacodynamic (PD) biomarker for DNA-PKcs activity, in all studied cell lines. As previously reported [Zenke et al., 2020], this IR-induced Ser2056 autophosphorylation was substantially impaired when the cells were exposed to 300nM of the DNA-PKcs inhibitor peposertib. As to the p53 status of the studied cell lines, total and phosphorylated p53 protein is absent in HPV+ cells due to the physiological ability of the viral E6 protein to degrade p53 [Hoppe-Seyler, 2018]. Similarly, the presence of the total but absence of the phosphorylated p53 protein in p53-mutated cell lines suggests that the loss of p53 function drives a phenotypic resemblance of these cells to HPV+ HNSCC lines.

Compared to exposure to IR alone, administration of the selective DNA-PKcs inhibitor peposertib before IR-induced DNA damage resulted in decreased cell viability and proliferation in all studied HNSCC cell lines. Our findings show that HPV+ cell lines and their derived tumors are more sensitive to IR, especially compared to p53 wild-type cell lines, and these observations are in line with published preclinical and clinical data [Nagel et al., 2013; Arenz et al., 2014]. We did not observe any IR dose-dependent responses in p53-mutated UM-SCC-10B and UM-SCC-14A cells combined with peposertib (their proliferation after 2 and 4Gy IR was affected to the same extent), likely due to the high IR sensitivity of p53-mutated cell lines [Anbalagan et al., 2021; Stewart-Ornstein et al., 2021]. This finding indicates that the same effect can be achieved by the addition of peposertib to a lower IR dose compared to IR alone.

Based on most of the readouts, all cell lines abrogate proliferation after combined treatment, but only p53-depleted cell lines predominantly arrest in the G2 phase of the cell cycle. As p53 is mainly involved in the transition from G1- to S-phase, p53-proficient cells cannot progress further in the cell cycle, while cells lacking p53 can pass the checkpoint but arrest in the G2-phase through p53-independent regulators [Kuntz et al., 2009; Kishi et al., 2001]. Partially in accordance with our findings, Sun et al. demonstrated that a single dose of 5Gy IR causes G1- and G2-M-phase arrests in p53 wild-type A375 melanoma and A549 lung cancer cell lines [Sun, 2019]. In their study, the combination of 5Gy IR with 1uM peposertib arrested the cells predominantly in the G2-M phase with no cells found in the S phase for three consecutive days. On the contrary, only partial and unstable arrest after combination treatment was observed in HPV+ HeLa cells.

Following DNA-PKcs inhibition, the IR-induced DNA damage cannot be repaired via NHEJ, and the cells presumably rely on alternative pathways including HR-based repair [Saleh-Gohari, N., & Helleday, T. et al., 2004; Zhao et al., 2017]. As our data show, the level of DNA damage induced by the combined treatment is significantly high even 48h post-treatment regardless of the cell line, confirming thus the persistent damage and delay in damage resolution causing activation of cell death/senescence pathways.

Our results further support the notion that cell fate after the induced DNA damage largely depends on its p53 status. By other studies, cells lacking functional p53 due to its degradation by HPV proteins or due to the presence of a loss-of-function mutation arrest upon the combined treatment in the G2 cell cycle phase and are subsequently eliminated by apoptosis. On the other hand, HNSCCs with functional p53 undergo senescence [Sun et al, 2019; Hannah et al, 2019; Wang et al, 2021; Carr et al, 2022; Ding et al, 2022]. High level of DNA damage induced by IR and inhibition of the NHEJ in the cells with proficient p53 strongly activates the p53/ATM pathway (p21, CHK1, CHK2 cell cycle proteins) and therefore induces premature senescence to protect genome integrity [Finzel et al, 2016; Sun et al, 2019]. In the absence of p53, cells suffer from the increased level of DNA double-strand breaks and die by mitotic catastrophe due to of severe chromosomal aberrations [Haines et al, 2021].

Although we could not observe the induction of apoptosis in p53 wild-type cell lines, cell death was detected after combination treatment in all cell lines, indicating that non-apoptotic cell death mechanisms such as necrosis might be involved. Importantly, we show that upon the combined peposertib and IR treatment, p53 wild-type tumor cells mainly undergo senescence, a non-proliferating cell state with a dual anti- and pro-tumorigenic phenotype. It has been demonstrated that other genotoxic chemotherapies induce senescent cells that can potentially create local and systemic inflammatory responses [Wang, 2022]. The reported in-vivo data indicates an early tumor regrowth of p53 wild-type xenografts upon initial response. Thus, although peposertib radiosensitizes HNSCC models regardless of their p53 and HPV status, a potentially cytotoxic and durable benefit from the combined treatment is selectively seen in HPV+ or p53-mutated tumors. If further studies would indicate that p53 wild-type HNSCC cancers do not benefit from the IR+peposertib combination, the treatment-induced senescent cells could be targeted with senolytic agents to eliminate the senescent cells following such a treatment combination [Carr et al., abstract, 2022].

Our in-vitro data are supplemented by in-vivo studies evaluating responses of either p53 wild-type or HPV+ HNSCCs to the combined IR+peposertib treatment. Radiation therapy was delivered by the SmART precision radiotherapy platform, which allows to image the lesion and to plan the radiation procedure similarly to in the clinic. Precision radiation targets the region of interest by sparing the healthy tissues from unnecessary irradiation. To the best of our knowledge, in previous publications assessing peposertib efficacy in combination with IR, cabinet irradiators were used and this is the first study to use precise IR treatment for this purpose. The significant suppression of tumor growth upon combination treatment in both p53 wild-type, as well as HPV+ models, is in line with recent data reported by Zenke et al [Zenke et al, 2020]. As to the tumor regrowth after treatment, we did not detect any recurrences of HPV+ tumors within the 45-day follow-up period, whereas p53 wild-type xenografts relapsed 30 days after treatment. Notably, Zenke et al. reported tumor regrowth in the p53-deficient HNSCC cell line FaDu 30 days after treatment initiation. However, a remarkable treatment response was achieved by increasing either the peposertib dose (up to 150mg/kg) or by applying a longer, 6-week radiotherapy regime with peposertib doses of up to 50mg/kg) [Zenke et al, 2020].

High levels of proliferating and apoptotic cells and low levels of senescent cells detected in HPV+ compared to HPV- can be one of the reasons for determining a better treatment response in HPV+ tumors. The vast difference in β-gal staining between peposertib and IR+peposertib treatment groups can be related to the differences in the tumor collection days counted from the treatment initiation (on average, 10.71±4.7 days for the peposertib group and 43.42±3.6 days for IR+peposertib groups).

In conclusion, diverse patterns of viability and cell cycle distribution upon the combined IR+peposertib treatment in HNSCC cells with dysfunctional p53 compared to p53 wild-type cells revealed that the outcome of such treatment depends on the p53 and the HPV status of a certain tumor. Although peposertib seems to radiosensitize HNSCC tumors independently of their genetic background, a better treatment response was observed both in-vitro and in-vivo in models with dysfunctional p53. HPV and p53 status thus emerge as critical factors that may shape treatment outcomes for HNSCC patients that could receive a combined IR and DNA-PKcs targeting treatment. Precise patient stratification based on these two parameters might be relevant also in the case of other malignant conditions. Pending further studies, whereas the treatment-induced senescence detected here in p53 wild-type models can become a target for the follow-up senolytic approach, the strong induction of apoptosis in p53 dysfunctional tumors could be synergistic with current pro-apoptotic therapies that are on the horizon in the clinic.

## Acknowledgments

We would like to thank Prof. Dr. D. Stroka and Dr. J. Gavini (both Department for BioMedical Research (DBMR), Visceral Surgery, University of Bern) for their valuable inputs for the in-vivo study, R. Riedo (DBMR, Radiation Oncology, University of Bern) for the assistance with experiments and L. Vassilev (formerly of EMD Serono, Billerica, MA, USA) for his valuable scientific input.

## Author contributions

Mi.M. and Y.Z. concepted, designed, and supervised the study. Li.H. and S.M.R. acquired, analyzed, and visualized experimental data. Lu.H. and A.Q. performed and analyzed the experiments. J.O. and A.Q. assisted with the SmART treatment planning. Ma.M. assisted with statistical analysis and data interpretation. J.A., D.M.A., Mi.M., and Y.Z. provided resources. Li.H., S.M.R., Y.Z., and Mi.M. drafted the manuscript and D.M.A. and J.A. revised the manuscript for important scientific content. All authors reviewed and approved the final version of the manuscript.

## Data availability

The data that support the findings of this study are available on a reasonable request from the corresponding author.

