## Supplemental figures for "HPV and p53 status as precision determinants of head and neck cancer response to DNA-PKcs inhibition in combination with irradiation"

### Supplementary Figures and legends

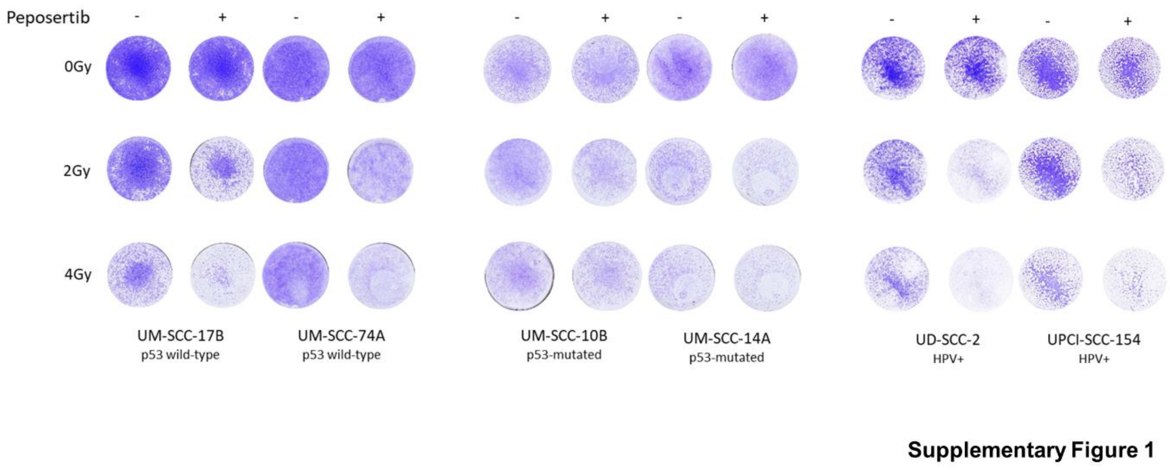


#### Supplementary figure 1.

Representative images of crystal violet assay.

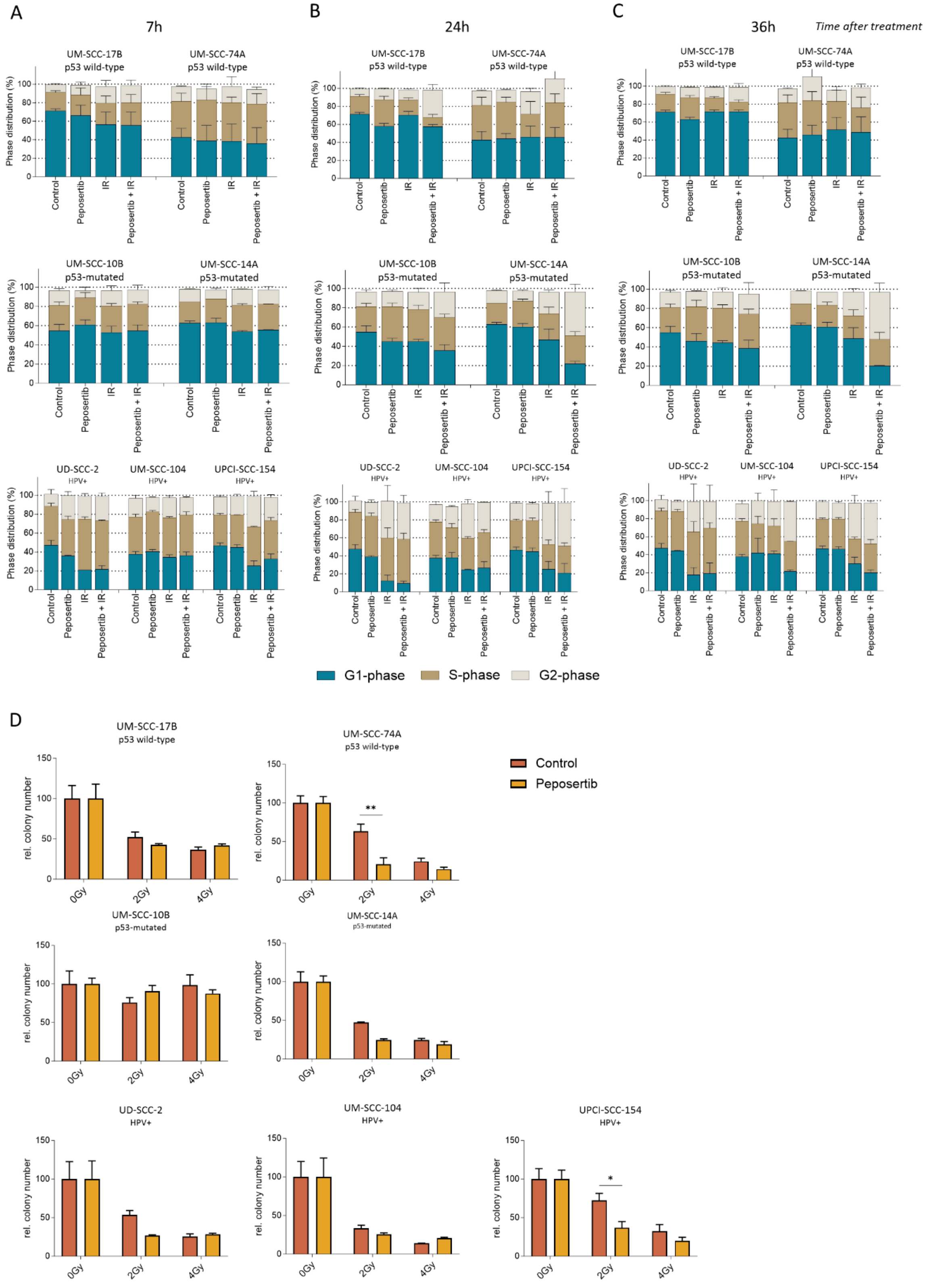


**Supplementary Figure 2**

#### Supplementary figure 2.

Cell cycle distribution data at 7h **(A)**, 24h **(B),** and 36h **(C)** after treatment *(p values are not shown)*. **(D)** Colony-forming assay (t*wo-way ANOVA with post hoc multiple comparisons. Data is represented as mean±SEM;* $p<0.05$, ** $p<0.01$, *** $p<0.001$, **** $p<0.0001$).

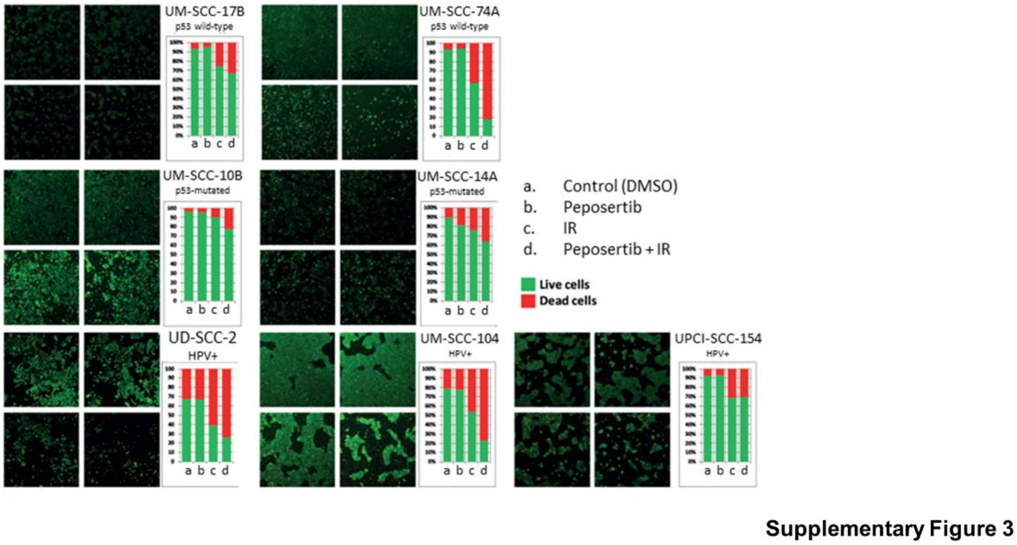


#### Supplementary figure 3.

Change of live-to-dead ratio observed in all cell lines. The LiveDead assay was performed 5 days after treatment.

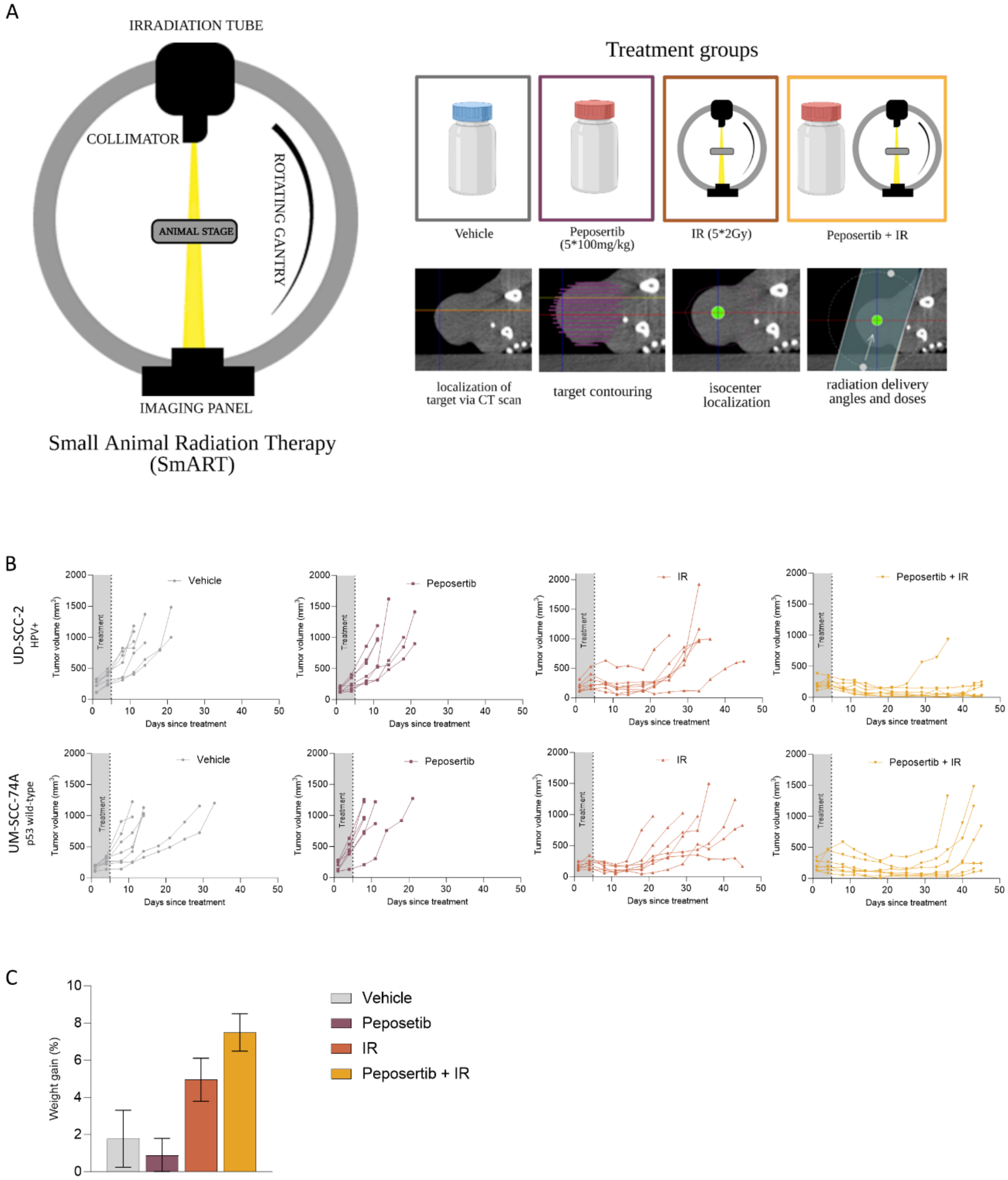


**Supplementary Figure 4**

#### Supplementary figure 4.

**(A)** In-vivo study treatment setup. **(B)** Individual tumor growth for each mouse (n=7) in all experimental and treatment groups. **(C)** Average weight gain of mice. **(D)** Xenograft tumor slices of UD-SCC-2 and UM-SCC-74A cell lines stained for proliferation and apoptosis show opposite staining patterns.
